# Context-specific and context-invariant computations of interval timing

**DOI:** 10.1101/2023.04.28.538792

**Authors:** Ahmad Pourmohammadi, Mehdi Sanayei

**Affiliations:** School of Cognitive Sciences, Institute for Research in Fundamental Sciences (IPM), Tehran, Iran; Center for Translational Neuroscience (CTN), Isfahan University of Medical Sciences, Isfahan, Iran

## Abstract

An accurate sense of time is crucial in flexible sensorimotor control and other cognitive functions. However, it remains unknown how multiple timing computations in different contexts interact to shape our behavior. We asked humans to perform timing tasks that differed in the sensorimotor domain (sensory timing vs. motor timing) and effector (hand vs. saccadic eye movement). To understand how these different behavioral contexts contribute to timing behavior, we applied a three-stage Bayesian model to behavioral data. We found that these behavioral contexts affect different stages of computations about time. Moreover, our results indicated that the mode of response also affects computations related to measuring and sensing time. These findings suggest that both context-specific and context-invariant computations contribute to shaping our timing behavior.

## Introduction

How the brain perceives time has been the focus of many studies over the past decades ^1^. The computational goal for measuring time at the present moment is to accurately and precisely track elapsed time within an ongoing interval. This computation is integral to sensorimotor control and decision-making, yet our subjective experience of time can be biased in different contexts. A classic example of this contextual calibration is Vierordt’s law, also known as the central tendency effect ^2, 3^. Vierordt showed that short temporal durations tend to be overestimated in a temporal reproduction task, whereas long durations tend to be underestimated ^3^. However, the underlying mechanism of these observations remains unexplained for more than a century. The central tendency effect is related to another feature of interval timing: Weber’s law. According to Weber’s law, the variability of temporal performance increases with the mean of the time interval, also known as scalar variability ^4-7^. Although information processing models based on the internal clock, memory-mixing, and internal noise were applied to explain contextual calibration and Weber’s law (for review see ^7, 8^), these approaches did not provide a quantitative prediction of the factors that contribute to contextual calibration. Furthermore, how the nervous system uses contextual calibration to improve timing behavior remains unknown.

To address the effects of contextual calibration on interval timing, a prior study used a Bayesian framework. Jazayeri & Shadlen ^9^ developed and compared three probabilistic observer models with different strategies (i.e. maximum-likelihood estimation, maximum a posteriori, and Bayes least-squares). They showed that observers used the Bayes least-squares strategy to reproduce temporal intervals. This finding was concluded based on the success of the Bayes least-squares model which provided an accurate description of behavioral data in a temporal reproduction task. The Bayesian framework suggests that the observer uses two sources of information to estimate a sample interval: sensory measurements (i.e., likelihood function) and the prior knowledge of the statistical distribution. This approach can also provide important insights into other aspects of temporal processing.

To understand the underlying mechanisms of timing, previous studies compared timing behavior in different contexts. Several psychophysical studies showed that timing behavior, especially scalar variability, is similar for different explicit timing tasks or effectors ^7, 10-14^. These results may suggest that the brain has a specialized set of circuits for measuring time across sensorimotor domains and effectors ^15^. However, electrophysiological and computational studies showed that time can be encoded through changes in neural population activity over time, or population state dynamics ^16-20^. Although dedicated and intrinsic models of timing, are not mutually exclusive ^21-24^, converging data from psychophysical, neuroimaging and electrophysiological studies supported partially overlapping distributed timing mechanisms for interval timing (for a review see ^1, 25, 26^). Indeed, A longstanding question is how these distributed timing circuits interact to shape temporal behavior. Also, it remains unknown how sensorimotor domain or motor response type affect computations about time. To address these questions, we applied and compared Bayesian models in different temporal contexts. Interval timing tasks in this study differed in either the sensorimotor domain (sensory timing vs. motor timing) and effector (hand vs. saccadic eye movement). Sensory timing refers to tasks in which decisions are based on the temporal structure of events while motor timing refers to tasks in which subjects time their own action. We hypothesized that the different behavioral contexts affect different stages of computations about measuring time.

## Result

In this study, we asked how sensorimotor domain or effector affects computations about time. To answer this question, first we compared behavioral data between different sensorimotor domains and different effectors.

Human subjects were asked to perform two different timing tasks and report their choices via a button press or eye movement (Fig. 1). In the reproduction task, subjects had to measure and reproduce sample intervals, t_s_. t_s_ were demarcated by two visual flashes and were selected from a discrete uniform distribution. The corresponding reproduction time was measured from the end of t_s_ to when subjects responded via a manual key press or a saccadic eye movement, in different blocks. In the discrimination task, subjects had to measure and compare two different sample intervals, t_s1_ and t_s2_, demarcated by three visual flashes. The values of t_s1_ were selected from a discrete uniform distribution, identical to the value of t_s_ in the reproduction task, and the values of t_s2_ were chosen from the distribution of t_s1_ ± [6, 12, 24, 48] % of t_s1_, randomly on each trial. Subjects compared t_s2_ with t_s1_ and chose whether t_s2_ was longer or shorter than t_s1_ via a manual key press or a saccadic eye movement. The two modes of response were used in different behavioral testing blocks. In both tasks subjects received visual feedback after each response.

**Figure 1.**
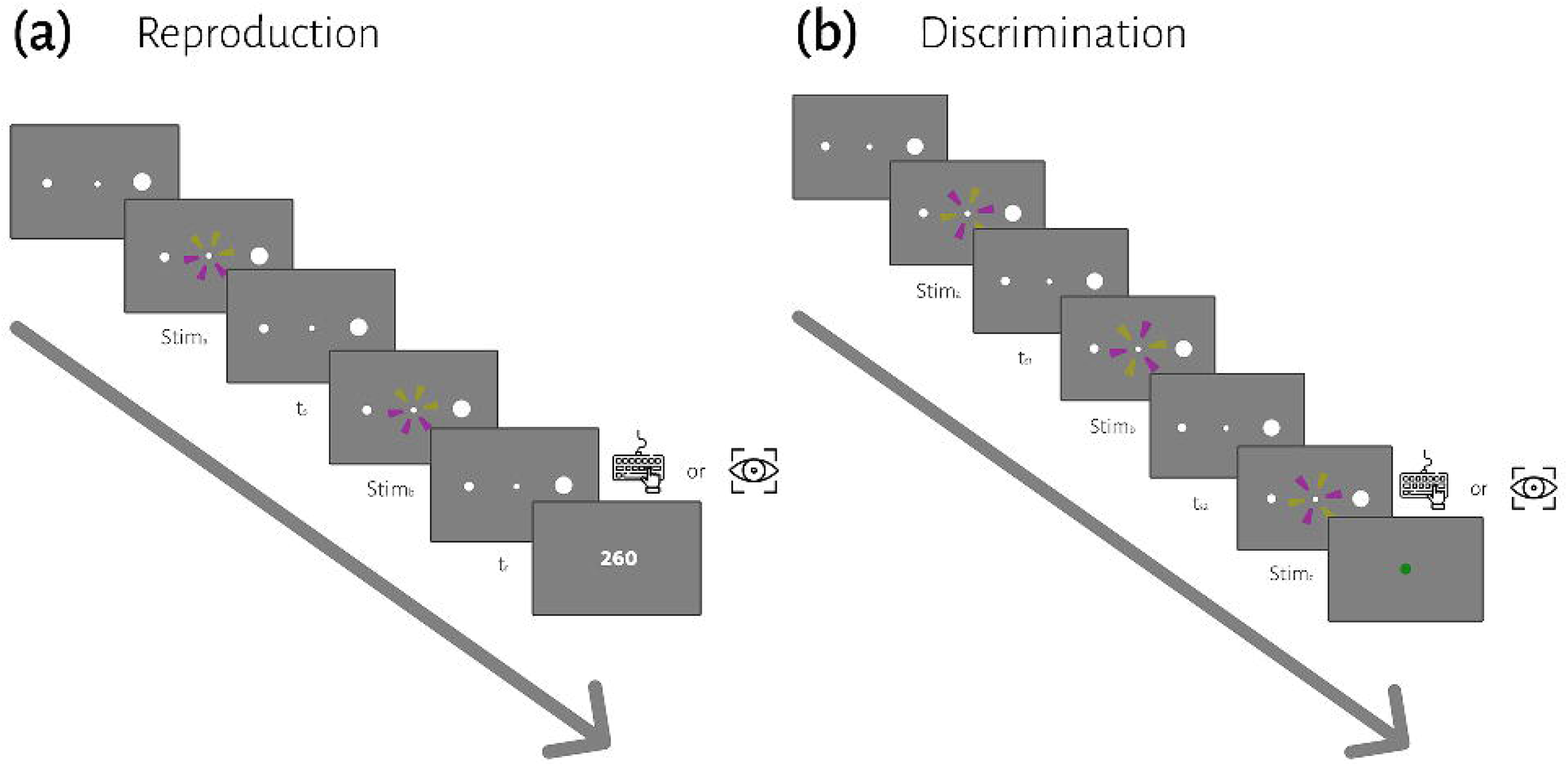
The sequence of trial events for the reproduction (a) and discrimination (b) tasks. (a) In the reproduction task, subjects had to measure and reproduce sample intervals, t_s_. After the subject acquired fixation and a random delay, t_s_ was demarcated by two visual flashes (stim_a_ and stim_b_). At the end of each trial, we showed the response error (t_r_ - t_s_) as feedback. (b) In the discrimination task, subjects had to measure and to compare two different sample intervals, t_s1_ and t_s2_, demarcated by three visual flashes (stim_a_, stim_b_, and stim_c_). t_s1_ were selected from a discrete uniform distribution, same as the t_s_ in the reproduction task. Subjects compared t_s2_ with t_s1_ and chose whether t_s2_ was longer or shorter than t_s1_ via a manual key press or saccadic eye movement in different blocks. At the end of each trial, visual feedback was presented to the subject.

Subjects showed both characteristic features of interval timing: central tendency bias and scalar variability (Fig. 2). First, response intervals (t_r_, mean of reproduction time and point of subjective equality for reproduction and discrimination, respectively) were systematically biased toward the mean of the presented distribution, a phenomenon known as central tendency bias. We quantified this central tendency bias with two parameters of a regression analysis (see Methods): the magnitude of the compressive bias and the indifference point. Second, the measurement of longer sample intervals engenders more uncertainty, a phenomenon known as scalar variability, as can be seen in increasing the standard deviation (σ) as interval time increased. Scalar variability was quantified with two parameters of Weber’s law (see Method), the time-dependent (Weber’s fraction) and time-independent parameters of Weber’s law.

**Figure 2.**
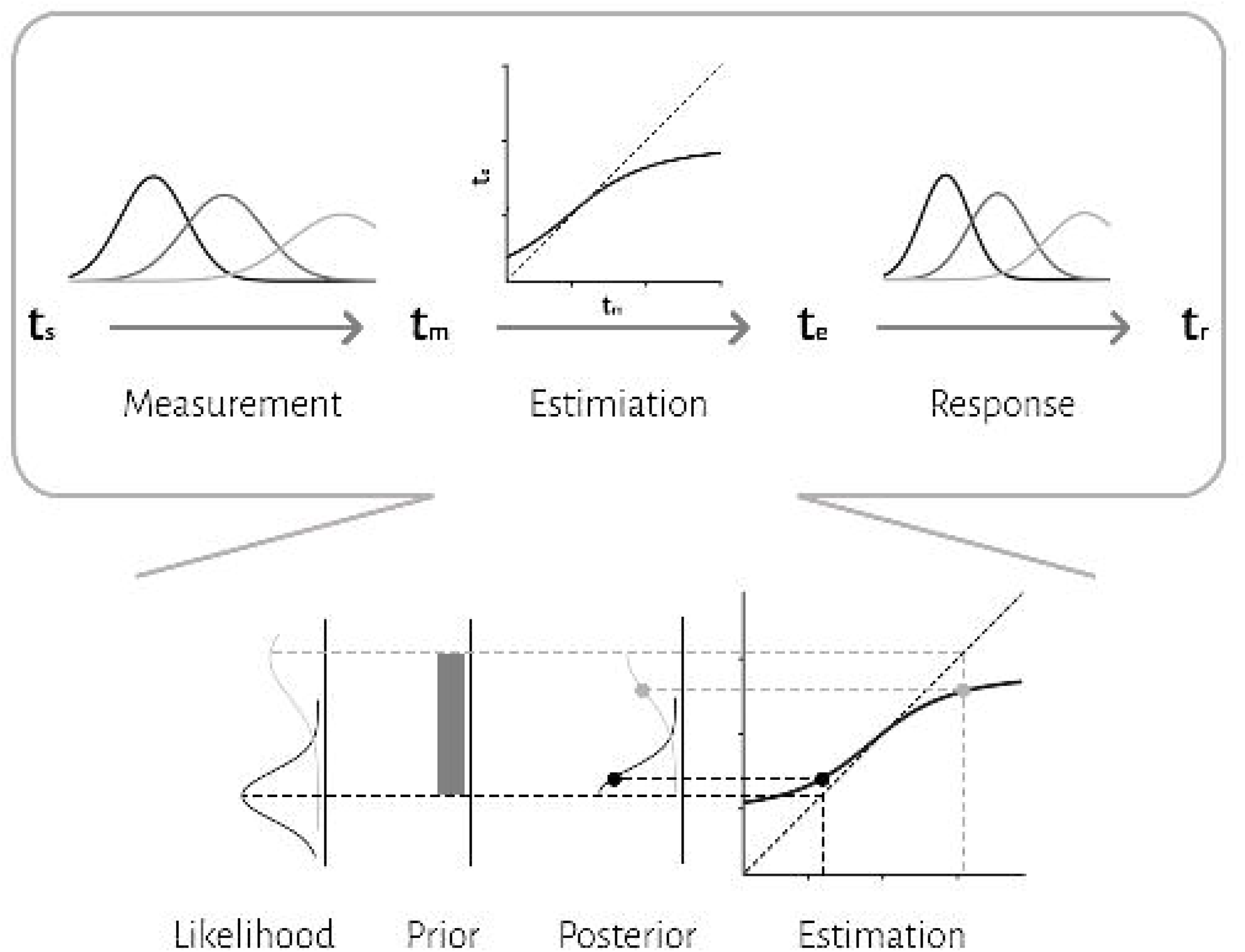
Timing behavior in the reproduction (a and c) and discrimination (b and d) tasks. (a) The distribution of reproduction times. (b) The proportion of long responses over conditions (t_s2_) for each interval (t_s1_). (c) Mean reproduction time as a function of interval duration in the reproduction task (filled symbols). Error bars show standard deviation of reproduction time. Solid lines show best-fitting linear regression function, whereas the dotted diagonal lines denote unbiased performance. Open symbol represents the estimated indifference point along with bootstrapped 95% confidence intervals. (d) Point of subjective equality (PSE) as a function of interval duration in the discrimination task. Error bars show standard deviation of fitted psychometric function.

To examine the effects of sensorimotor domain and effector on temporal processing, we compared central tendency bias and scalar variability between different sensorimotor domains and effectors. We did not find any significant difference between blocks employing key presses or saccadic eye movements when we compared compression bias (F = 2.62, p = .11), indifference point (F = .55, p = .44), Weber’s fraction (F = 0.00, p = .99) and time-independent parameter (F = 3.14, p = .09). Similarly, we did not find any significant difference between reproduction and discrimination blocks when we compared compression bias (F = 1.78, p = .19), indifference point (F = .35, p = .58), Weber’s fraction (F = 1.66, p = .21) and time-independent parameter (F = 2.10, p = .16). The interaction of effector and task on the indifference point was significant (F = 12.32, p < .01) but further analysis using HSD Tukey test did not find any significant effect in multiple comparison between groups (Table S1).

### The Bayesian observer model

To further evaluate the effects of sensorimotor domain and effector on temporal processing and to understand the computations from which these effects might arise, we applied a Bayesian model with three stages, measurement, estimation and motor response (Methods, Fig. 3). In the first stage, an observer takes noisy measurements, t_m_, from sample intervals, t_s._ Measurement noise was modeled as a Gaussian function. The second stage is a Bayes least-squares (BLS) estimator. The estimator is a deterministic function, f(t_m_), that maps t_m_ to t_e_. In the third stage, the Bayesian observer uses t_e_ to respond, t_r_. The relationship between t_r_ and t_e_ is characterized by motor noise, which was modeled by a Gaussian distribution. The measurement constant coefficient of variation, w_m_, is the model’s free parameter for measurement stage (the first stage) and the response constant coefficient of variation, w_r_, is the model’s free parameter for motor response stage (the third stage). The estimation stage (the second stage) does not invoke any additional free parameters.

**Figure 3.**
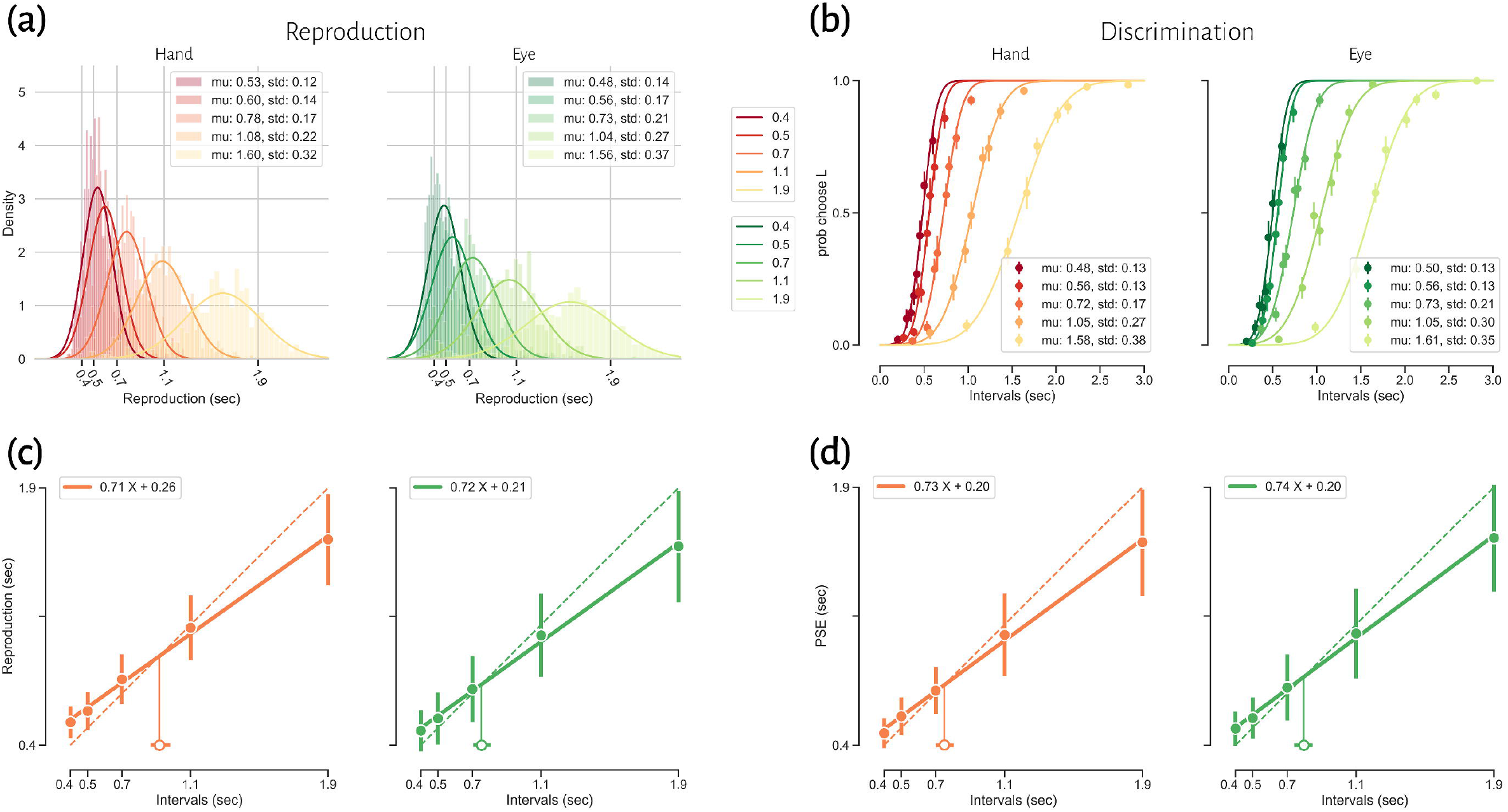
The three-stage architecture of the Bayesian observer model. In the first stage, the observer takes noisy measurements, t_m_, from sample intervals, t_s._ Measurement noise was modeled as a Gaussian function. The second stage is a Bayes least-squares (BLS) estimator. The estimator is a deterministic function, f(t_m_), that maps t_m_ to t_e_. In the third stage, the observer uses t_e_ to respond, t_r_. The relationship between t_r_ and t_e_ is characterized by motor noise, which was modeled by a Gaussian distribution. An illustration of how compressive biases arise in the model is depicted at the bottom. According to the Bayesian approach, an observer combines noisy sensory measurements (i.e., likelihood) with the prior knowledge of the statistical distribution of the stimulus to improve behavior.

We fitted the parameters of the Bayesian model, w_m_ and w_r_, for each subject on the basis of t_r_. Then we compared w_m_ and w_r_ between different tasks and different motor response types. Both effector (F = 9.38, p < .01) and tasks (F = 5.02, p < .03) had significant effects on w_m_, but the interaction was not significant (F = 1.83, p = .19). Effector had a significant effect on w_r_ (F = 8.97, p < .01), while task did not (F = 3.09, p = .09). Their interaction though reached significant level (F = 6.60, p = 0.02). Additional analysis showed that w_r_ was significantly larger in the discrimination with eye (0.28 ± 0.09) than in the reproduction with hand (0.22 ± 0.05, adjusted p = .04, Table S1).

To compare subjects’ responses to those predicted by the model, we simulated each subject’s responses using the fitted Bayesian model and compared model predictions to the actual responses using the bias and variability statistics. The correlations showed that our Bayesian model can predict subjects’ responses in each effector and each task (rs > 0.50 and ps < .01, Fig. 4).

**Figure 4.**
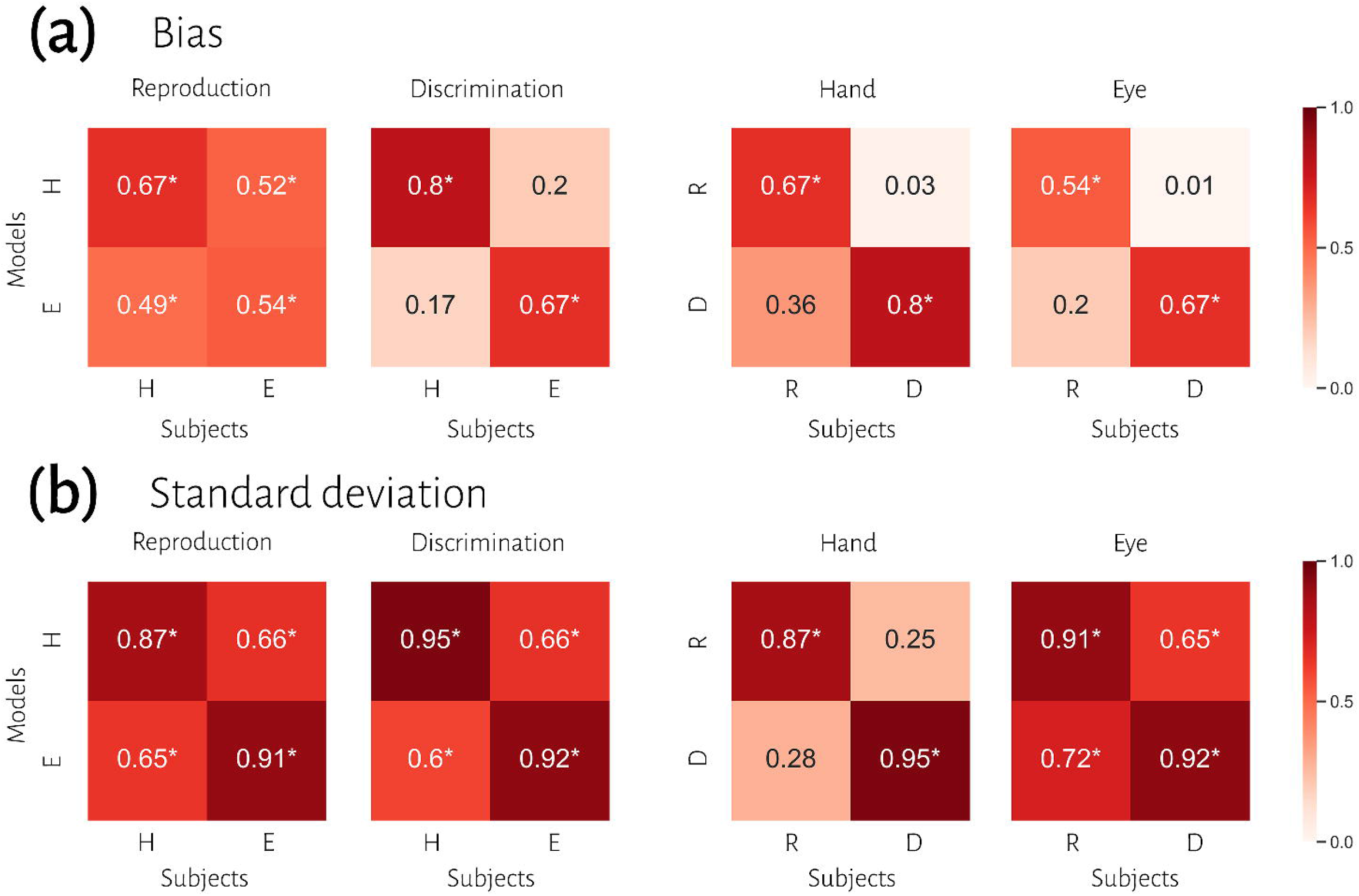
Comparison of timing behavior in human and model observers. (a) The correlation between subject’s bias and simulated data from best fitting model for each subject. (b) The correlation between subject’s variability and simulated data from best fitted model for each subject. Abbreviations: reproduction (R), discrimination (D), hand (H), eye (E). *: denotes a significant correlation

To evaluate the effect of effector, we compared subjects’ responses in blocks using a key press to model predictions in blocks using a saccadic eye movement and vice versa in the same tasks. In reproduction, bias was significantly correlated between subjects’ responses in each effector and model fitted to the other effector (model hand, data eye: r = .52, p < .01; model eye, data hand: r = .49, p < .01). However, in the discrimination task, bias was not significantly correlated between subjects’ responses and model predictions with different effectors (Fig. 4). In contrast, variability was significantly correlated between subjects’ responses and model predictions with different effectors in both tasks (rs > .60, ps < .01).

To evaluate the effect of tasks, we compared subjects’ responses in the reproduction task to model predictions in the discrimination task and vice versa with the same effector. Bias was not significantly correlated between subjects’ responses in each task and model fitted to the other task (Fig. 4). Variability was significantly correlated between subjects’ responses and model predictions with different task in the eye blocks (model reproduction, data discrimination: r = .65, p < .01; model discrimination, data reproduction: r = .72, p < .01) but not in hand blocks (Fig. 4). The scatter plot for each correlation analysis is shown in Fig. S1.

## Discussion

In this study, we answered how behavioral context affect computations about time. We found that temporal accuracy (i.e., compression bias and indifference point) and precision (i.e., Weber’s parameters) were not different between these behavioral contexts. Previous studies also reported that temporal precision is correlated across different sensorimotor domains ^27^ and different effectors ^11^. Although these observations may qualitatively suggest that subjects who are good timers in one behavioral context are also good timers in another one, converging data from psychophysical studies remain controversial (for review see ^28^). To evaluate and predict the contribution of behavioral contexts on timing computations, we applied the Bayesian framework to our behavioral data.

Bayesian models have made great advances in describing a variety of cognitive functions including interval timing ^29, 30^. This approach proposes that an optimal observer combines noisy sensory measurements with the prior knowledge of the statistical distribution of the stimulus to improve behavior. This behavioral computation is interconnected with the familiar trade-off between accuracy and precision: the prior-dependent bias increases for less reliable measurements. Previous studies used Bayesian models to elucidate how prior knowledge is formed in different behavioral contexts ^31-34^. They suggested that representation of prior knowledge is sensory specific after extended training. Roach et al. ^35^ extended this approach to interval timing. In a temporal reproduction task, they applied a Bayesian model to study how priors are learned and how they are generalized to different behavioral contexts. They showed that participants formed a single prior by generalizing across intervals coupled with different sensory modalities in early sessions. However, prior generalization was not occurred and participants formed multiple, and separate, priors across duration distributions when coupled with different effectors. The authors suggested that representation of prior knowledge is effector specific, but not sensory specific, in the temporal reproduction task without extended training. However, it remained unclear how behavioral context contribute to timing computations.

Motivated by this question and the success of previous Bayesian models of interval timing ^9, 35, 36^, we applied a Bayesian model with three stages to our timing data (Fig. 3). In this model, the first two stages contribute to bias, and the third stage contributes to response variability. Response variability is also affected by the posterior uncertainty. We showed that the Bayesian model can accurately describe behavioral data in different sensorimotor domains and different effectors. Previous studies also reported that Bayesian models with the Bayes least-squares strategy in the estimation stage described behavioral data in temporal reproduction task with hand response ^9, 36^. Our results supported these studies and extended these findings to sensory timing and saccadic eye movement.

Our results demonstrate that the Bayesian model for each task could not describe bias in other task in either hand or eye blocks. The first two stages of Bayesian model contribute to response bias. The lack of correlation and significantly different measurement noise (w_m_) between different tasks suggest that timing computation in the measurement stage is different between sensorimotor domains. In other words, the measurement stage is sensorimotor domain specific which suggests that the brain might use different mechanisms or computations for timing when it knows that it has to engage in sensory vs. motor timing tasks. We also showed that the Bayesian model for each task can accurately describe variability in the other task in eye blocks, but not in hand blocks. This finding suggests that although motor noise (w_r_) is not significantly different between tasks, timing computation in the motor stage is not correlated between sensorimotor tasks in hand blocks.

An interesting finding we made is that the Bayesian model for each effector can predict bias in other effector in the reproduction task, but not in the discrimination task. The lack of correlation and significantly different measurement noise (w_m_) between effectors suggest that the measurement stage is also effector specific in sensory timing, but not in motor timing. We also showed that the Bayesian model for each effector could predict variability in other effector in the reproduction or discrimination task. We showed that although timing computation in the motor stage was significantly correlated between effectors, w_r_ was significantly higher in the eye compared to hand blocks.

In sensory and motor timing, the observer’s task can be explained as accumulating evidence in the time domain and comparing it to a bound, similar to what has been shown in decision making ^37^. This evolving decision variable would then lead to a motor action when reaches a bound. In our sensory timing task, temporal decisions and motor responses are made by choosing between two alternatives. However, in the motor timing task, decisions and motor responses are made by choosing when to act. De Lafuente et al., ^38^ used a two-alternative forced choice (2-AFC) random dots motion discrimination task showed that when decisions are communicated by different effectors neurons in the medial intraparietal (MIP) area exhibit different firing activity. Instead, decision-related activity was observed in the lateral intraparietal (LIP) area in both hand and eye blocks. We argue that at the computational level, when a temporal decision (or maybe any other decision) has to be made in a 2-AFC task, temporal measurement (the first stage in our model) raises from different computations in eye vs. hand blocks. But when an observer wants to decide when to act, a 1-AFC task, temporal measurement had the same computations between different effectors.

The current study suggests overlapping and distributed computations as the underlying mechanism of timing in different contexts. Previous neuroimaging and electrophysiological studies also support this idea at the implementational level of analysis (for review see ^1, 25, 26^). Several studies showed that depending on the task and timescale, many areas had been implicated in different temporal contexts ^39-46^. The results of these previous studies suggest that time is encoded in both context invariant and context specific areas. In this framework, context specific time representation can be encoded through changes in neural population activity over time in distributed neural circuits ^1, 25^. However, a more detailed understanding of the neural substrates of context invariant computations is lacking. These computations might be implemented in overlapping neural circuits. The effects of neuromodulatory systems on distributed neural circuits can also play a critical role in context invariant computations of time. Previous studies reported that the basal ganglia are activated in timing tasks with different effectors ^42^, sensorimotor domains ^47, 48^, and duration scales ^49^. These studies suggested that context invariant computations might be implemented in the basal ganglia. Another study also reported that optogenetic manipulation of substantia nigra pars compacta (SNc) dopamine neurons can modify timing behavior ^50^. The results of this study suggested that dopaminergic projections from the SNc to the striatum could modify striatal population dynamics. Considering that the basal ganglia is interconnected with widespread regions of the cerebral cortex and subcortical areas, it is possible that basal ganglia can modify distributed neural circuits in timing tasks. Future studies are needed to address this question and to understand how these different timing computations are implemented in the brain.

## Methods

### Subjects

We enrolled 41 healthy human subjects, (22 females, 26.76 ± 4.30 years old, reported as mean ± sd). All participants were informed of the general purpose of the study but were naïve about the scientific questions and tasks, except for two who were the authors. Subjects had normal or corrected-to-normal vision and no history of neurological or psychiatric disorders. Written informed consent was obtained from all participants before the start of the study. Subjects performed three psychophysics tasks, each task with two different motor response types in separate sessions. Here we only report results from two of those tasks. We enrolled the same subjects in all tasks, and the order of tasks was counterbalanced between subjects. The Ethics Committee of the Institute for Research in Fundamental Sciences (IPM) approved this study.

### Apparatus

The experiments were carried out on a computer running Linux operating system, on MATLAB (2016b), with Psychtoolbox 3 extension ^51-53^. Stimuli were presented on a monitor (17”) placed ∼57 cm from the subject with a 60 Hz refresh rate. The subject sat comfortably on a chair in a dimly lit room to participate in this study, with the head stabilized by a head and chin rest. An EyeLink 1000 infrared eye tracking system (SR Research, Mississauga, Ontario) was used to record eye movements at 1kHz.

### Temporal reproduction task

Each trial began with the presentation of a central fixation point (diameter: 0.2°) and two peripheral targets (left target: 0.5° diameters, right target: 2° diameter, 10° eccentricity). After the subject acquired fixation within a ± 1° of fixation point, a trial would start. After a random delay (500ms plus a random sample from an exponential distribution with a mean of 250ms), two similar wheel-like stimuli (2.5° diameter) were flashed (for 26.6ms each) sequentially around the fixation point. The presented stimuli were a circle consisting of 6 sectors of equal size. The sectors were colored yellow (150 150 50 in RGB space) and purple (150 50 150), three each. The subject measured the time between two flashes, sample interval or t_s_, and produced a matching interval, t_r_, by pressing the right arrow key or making a saccade to the right target, depending on the block. Across trials, t_s_ was sampled from one of 5 discrete values pseudorandomly (400, 500, 700, 1100, 1900ms, uniform distributions). At the end of each trial, we showed the response error (t_r_ - t_s_) as feedback for 0.8 sec to the subject. The inter-trial interval was 1.2 sec, and each block contained 40 trials. Each subject participated in 6 blocks. In 3 blocks, they responded with their hand, and in 3 blocks, they responded with a saccadic eye movement. The order of response type was counterbalanced between subjects.

### Temporal discrimination task

Each trial started with the presentation of a fixation point and two peripheral targets (the same as the reproduction task). After a random delay, the first interval (t_s1_) started with the presentation of the first stimulus and ended with the presentation of the second stimulus (Fig. 1b). The second interval (t_s2_) immediately started with the second stimulus and ended with the presentation of the third stimulus. The stimuli were the same as the ones we used in the reproduction task, but the order of yellow and purple sectors was different. During the t_s1_ or t_s2_ interval, only the fixation point was shown (i.e., empty interval). The subject had to compare t_s2_ with t_s1_; If t_s2_ was longer than t_s1,_ then they had to press the right arrow key or make a saccade to the right target, if t_s2_ was shorter than t_s1,_ then they had to press the left arrow key or made a saccade to the left target. Across trials, t_s1_ was sampled from the same 5 discrete values as in the reproduction task. Duration of t_s2_ was t_s1_ duration ± [6, 12, 24, 48]% of t_s1_ duration, selected pseudorandomly on each trial. Feedback was shown at the end of each trial for 0.8 sec (a green circle with a 1.25° diameter for correct trials and a red circle of the same size for incorrect trials). The inter-trial interval was 1.2 sec, and each block contained 40 trials. Each subject participated in 12 blocks, half with hand response and the other half with eye response. The order of response type was counterbalanced between subjects.

### Analysis of behavioral data

In the reproduction task, we excluded outlier t_r_ which identified with Interquartile range method (for each subject; 166 from 10080 trials in total). we plotted mean of reproduction time (t_r_) as a function of interval duration (t_s_) for each subject and fitted a linear regression function to them. The fitted regression line yielded the magnitude of the compressive bias (C = 1 - the slope of the regression ^35^) and the indifference point (IP, as the time value at which the fitted regression line intersects the diagonal unity line ^35^). We also evaluated the relationship between the interval duration (t_s1_^2^) and timing variance (σ^2^) by fitting another linear regression, according to Weber’s law. The resulting slope and intercept correspond to the time-dependent (Weber’s fraction) and time-independent processes, respectively. These analyses were done separately for blocks that subject responded by hand and saccadic eye movement.

In the discrimination task, psychometric curves were generated by plotting the proportion of long responses over conditions (t_s2_) for each interval (t_s1_). These points were then fitted by a Gaussian cumulative density function with the non-linear least squares method. The mean (μ) of the fitted curve provided the point of subjective equality (PSE; the time value at which subjects judged t_s2_ was equal to t_s1_). In order to evaluate how well the psychometric function fitted with data, we measured the goodness-of-fit. We excluded poorly fitted conditions (R ≤ .70). Subjects with more than two excluded conditions were excluded entirely from all analyses (4 subjects). We characterized the relationship between interval duration (t_s1_) and PSE by fitting a linear regression. The results of this analysis yielded C and IP, as described before. We also evaluated the relationship between the interval duration (t_s1_^2^) and variance (σ^2^) of the fitted Gaussian cumulative density function by fitting another linear regression to calculate Weber’s parameters. These analyses were performed separately for blocks in which subjects responded by hand or saccadic eye movement.

We excluded subjects if the calculated indifference point was outside the range of presented intervals (i.e., shorter than 400 and longer than 1900ms) in the reproduction task (4 subjects) or the discrimination task (4 subjects, one was shared with exclusion based on the goodness-of-fit). Based on these exclusion criteria, 11 subjects were excluded from all analyses, and 30 remained.

To examine whether these computed variables were different between sensorimotor domains and between effectors, for each variable, we performed a two-way repeated measure ANOVA. Wherever we found a significant interaction between sensorimotor domain and effector, we applied HSD Tukey statistical test to control for multiple comparisons. Statistical analysis was performed with Python 3.8.5. A significant level was defined as p-value less than 0.05.

### The Bayesian observer model

We applied a Bayesian model with three stages ^9^: measurement, estimation and motor response (Fig. 3). In the first stage, an observer takes noisy measurements, t_m_, from sample intervals, t_s._ Measurement noise was modeled as a Gaussian function. We modeled the measurement stage as a Gaussian distribution with mean t_s_ and standard deviation w_m_t_s_, whereby the standard deviation grows as a constant fraction (w_m_) of the mean. This stage is also known as the likelihood function, λ:

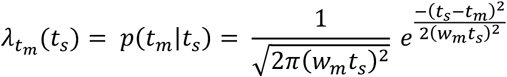

In the second stage, the Bayesian model combines the likelihood function and prior and use mean of the posterior to map the resulting posterior probability distribution onto an estimate, t_e_. Bayes least-squares (BLS) was used as the mapping rule in our model.

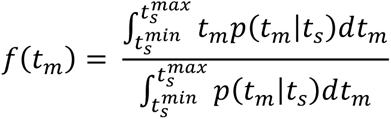

In the third stage, the ideal observer uses t_e_ to respond, t_r_. The relationship between t_r_ and t_e_ is characterized by motor noise, which was modeled by a Gaussian distribution with mean t_e_ and standard deviation w_r_t_e_.

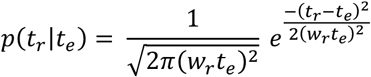

In the reproduction task, our psychophysical data consisted of pairs of sample intervals and reproduction times (t_s_ and t_r_). We derived a direct relationship between reproduction times and sample intervals in the Bayesian model ^9, 36^. This formulation was then used to describe subject’s responses in each effector, separately ^9, 36^. In the discrimination task, we transformed the psychometric function (Gaussian cumulative function) to response distribution (Gaussian probability function) with same parameters, as previous studies described ^11, 23, 54^. Responses were generated from this distribution and we had pairs of sample intervals and responses. Then same Bayesian formulation was used to describe subjects’ responses for each effector:

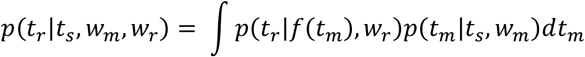

We maximized the likelihood of model parameters w_m_ and w_r_ across all t_s_ and t_r_ values. Maximum likelihood estimation was performed with minimize function in SciPy library, using the Nelder-Mead downhill simplex optimization method. We evaluated the success of the fitting procedure by repeating the search with several different initial values.

## Data availability

The data supporting the findings of this study are available from the corresponding author upon reasonable request.

## Acknowledgments

We are grateful to A. Sarabi-Jamab for advice on modeling and Y. El-Shamayleh for her feedback on the previous version of the manuscript.

