## Supplementary Information for "Context-specific and context-invariant computations of interval timing"

**Table S1.** Timing behavior between sensorimotor domains and between effectors. Values are reported as mean ± std.

Abbreviations: the magnitude of the compressive bias (C), indifference point (IP), Weber's fraction (Web_s), time-independent parameters of Weber's law (Web_i), measurement constant coefficient of variation ($w_m$), response constant coefficient of variation ($w_r$).

*: denotes a significant difference in two-way repeated measure ANOVA results.

| | Reproduction | | Discrimination | | ANOVA |
|---|---|---|---|---|---|
| | Hand | Eye | Hand | Eye | |
| C | 0.31 ± 0.13 | 0.31 ± 0.12 | 0.30 ± 0.14 | 0.25 ± 0.10 | effector: F = 2.85, p = 0.10<br>task: F = 1.78, p = 0.19<br>effector×task: F = 1.59, p = 0.22 |
| IP | 0.92 ± 0.24 | 0.75 ± 0.20 | 0.82 ± 0.26 | 0.92 ± 0.38 | effector: F = 0.55, p = 0.44<br>task: F = 0.35, p = 0.58<br>**effector×task: F = 12.32, p < 0.001*** |
| Web_s | 0.023 ± 0.023 | 0.031 ± 0.020 | 0.047 ± 0.090 | 0.040 ± 0.067 | effector: F = 0.00, p = 0.99<br>task: F = 1.66, p = 0.21<br>effector×task: F = 2.31, p = 0.14 |
| Web_i | 0.010 ± 0.011 | 0.021 ± 0.016 | 0.020 ± 0.033 | 0.025 ± 0.027 | effector: F = 3.14, p = 0.09<br>task: F = 2.10, p = 0.16<br>effector×task: F = 0.77, p = 0.39 |
| $w_m$ | 0.33 ± 0.13 | 0.27 ± 0.11 | 0.26 ± 0.11 | 0.24 ± 0.09 | **effector: F = 9.38, p < 0.01***<br>**task: F = 5.02, p = 0.03***<br>effector×task: F = 1.83, p = 0.19 |
| $w_r$ | 0.22 ± 0.05 | 0.28 ± 0.05 | 0.27 ± 0.12 | 0.28 ± 0.09 | **effector: F = 8.97, p = 0.01***<br>task: F = 3.09, p = 0.09<br>**effector×task: F = 6.60, p = 0.02*** |

**Figure S1.** Extended comparisons of timing behavior in human and model observers. (a) The correlation between subject's bias and simulated data from best fitted model for each subject. (b) The correlation between subject's variability and simulated data from best fitted model for each subject.

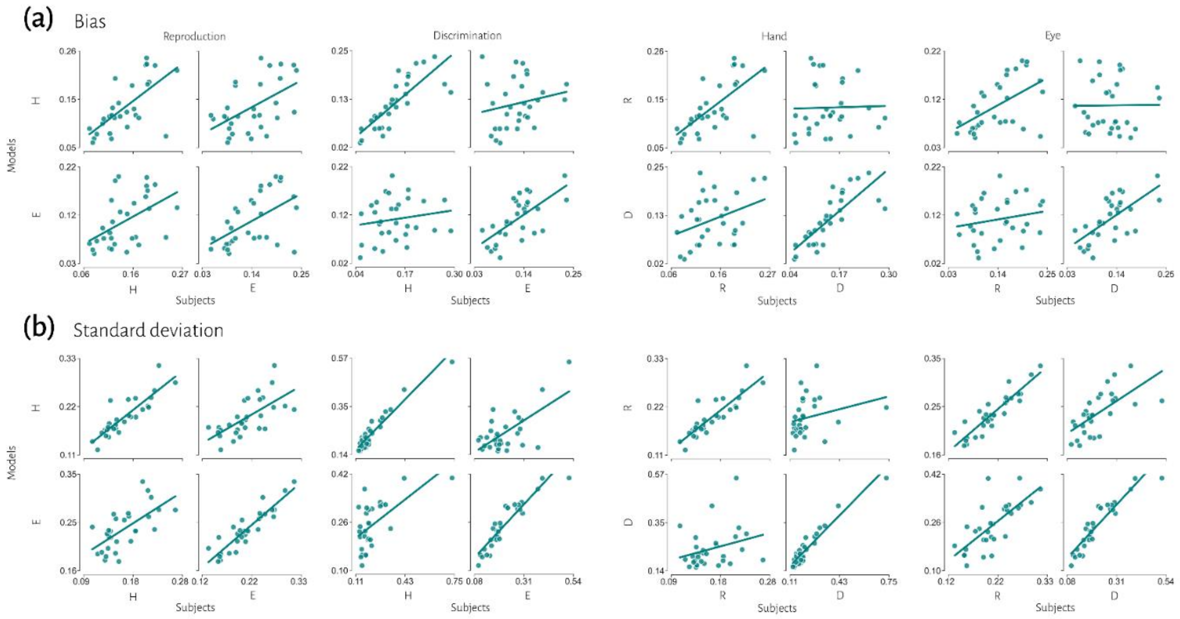
